# The genetic architecture of post-zygotic reproductive isolation between *Anopheles coluzzii* and *An. quadriannulatus*

**DOI:** 10.1101/2020.05.19.104786

**Authors:** Kevin C. Deitz, Willem Takken, Michel A. Slotman

## Abstract

The *Anopheles gambiae* complex is comprised of eight morphologically indistinguishable species and has emerged as a model system for the study of speciation genetics due to the rapid radiation of its member species over the past two million years. Male hybrids between most *An. gambiae* complex species pairs are sterile, and some genotype combinations in hybrid males cause inviability. We investigated the genetic basis of hybrid male inviability and sterility between *An. coluzzii* and *An. quadriannulatus* by measuring segregation distortion and performing a QTL analysis of sterility in a backcross population. Hybrid males were inviable if they inherited the *An. coluzzii* X chromosome and were homozygous at one or more loci in 18.9 Mb region of chromosome 3. The *An. coluzzii* X chromosome has a disproportionately large effect on hybrid sterility when introgressed into an *An. quadriannulatus* genetic background. Additionally, an epistatic interaction between the *An. coluzzii* X and a 1.12 Mb, pericentric region of the *An. quadriannulatus* 3L chromosome arm has a statistically significant contribution to the hybrid sterility phenotype. This same epistatic interaction occurs when the *An. coluzzii* X is introgressed into the genetic background of *An. arabiensis,* the sister species of *An. quadriannulatus*, suggesting that this may represent one of the first Dobzhansky–Muller incompatibilities to evolve early in the radiation of the *Anopheles gambiae* species complex. We describe the additive effects of each sterility QTL, epistatic interactions between them, and genes within QTL with protein functions related to mating behavior, reproduction, spermatogenesis, and microtubule morphogenesis, whose divergence may contribute to post-zygotic reproductive isolation between *An. coluzzii* and *An. quadriannulatus*.

## Introduction

The study of speciation genetics and genomics has largely focused on the evolution of hybrid dysfunction (Coyne and Orr, 2004; Dobzhansky, 1937) because it represents the first step in the formation of post-zygotic reproductive isolation between diverging populations. The evolution of post-zygotic isolation mechanisms is positively correlated with the time of divergence between two putative species, and is impacted by the number of mutations separating two allopatric lineages at any given time, the number of incompatibilities per mutation, and the fitness of each incompatibility (Orr and Turelli, 2001).

Post-zygotic isolation occurs as Dobzhansky-Müller incompatibilities (DMI) arise between populations evolving in allopatry (Dobzhansky, 1936; Müller, 1940). Upon secondary contact of genetically divergent populations, epistatic loci on parental chromosomes are unable to function effectively in hybrids. The resulting gene products in hybrids can have unpredictable, deleterious epistatic interactions (Maheshwari and Barbash, 2011). Orr (1995) and Orr and Turelli (2001) proposed the “snowball effect” of DMI evolution, which predicts that the number of DMIs accumulating between lineages “snowballs”, or increases faster than linearly (see Kalirad and Azevedo, 2017 for explicit testing of the underlying assumptions of this model). The snowball effect has been supported by empirical evidence of the rate of DMI evolution between *Drosophila* (Matute et al., 2010; Matute and Gavin-Smyth, 2014) and *Solanum* sibling species pairs (Moyle and Nakazato, 2010).

DMI can evolve as a result of various mechanisms in either an ecological speciation (divergent evolution due to selection pressures imposed by novel environments) or mutation-order speciation scenarios (fixation of different, incompatible mutations in allopatric populations experiencing the same environmental conditions) (Mani and Clarke, 1990; Nosil and Flaxman, 2011). These mechanisms include mutation-driven co-evolution (e.g. copper tolerance in *Mimulus guttatus*, MacNair, 1983; MacNair and Christie, 1983), gene duplication (e.g. the *OdsH* gene in *Drosophila simulans × D. mauritiania* backcross males, Ting et al., 2004; Sun et al., 2004), gene transposition (e.g. the *JYAlpha* gene in D*. melanogaster × D. simulans* backcross males, Masley et al., 2006), or a molecular arms race (e.g. divergent NB-LRR alleles in *Arabidopsis thaliana* strains, Bomblies et al., 2007; Bomblies and Weigal, 2007; Bakker et al., 2006). Furthermore, divergence in gene regulatory regions between parental species can result in in allelic imbalance in hybrids due to incompatible *cis-* and/or *trans*-acting factors (Graze et al., 2012), and segregation distortion of parental loci can skew sex ratios of hybrid progeny (Orr and Irving, 2005; Phadnis and Orr, 2009).

The *Anopheles* (*An.*) *gambiae* complex is comprised of eight morphologically indistinguishable species and has emerged as a model system for the study of speciation genetics due to the rapid radiation of its member species over the past two million years (Fontaine et al., 2015). All members of the *An. gambiae* complex are competent vectors of human malaria parasites, though they differ in their host specificity, ranging from highly anthropophilic (*An. gambiae s.s.* and *An. coluzzii*) (Takken and Knols, 1990; Dekker and Takken, 1998; Pates et al., 2001a) to almost entirely zoophilic (*An. quadriannulatus* A) (White, 1974; Gibson, 1996; Dekker and Takken, 1998; but see Pates et al., 2001b). From a vector control stand point, recent advances in the development of transgenic mosquito control methods have made understanding the permeability of species boundaries even more urgent (e.g. Kyrou et al., 2018) because it is important to know whether transgenic alleles have the potential to introgress between target and non-target species.

F1 hybrid female progeny of crosses between species of the *Anopheles gambiae* complex are fertile, but, with the exception of the *An. gambiae* - *An. coluzzii* species pair, males range from semi-to completely sterile (White, 1974; Hunt, 1998). A previous QTL analyses of sterility in bi-directional backcross hybrids identified regions in the *An. coluzzii* genome that cause male and female sterility when introgressed into an *An. arabiensis* genetic background, and vice versa (Slotman et al., 2004; Slotman et al., 2005). One or more loci on the X chromosome contribute to the effect of the X chromosome on the hybrid sterility phenotype in both directions of the cross. Unfortunately, recombination on the X chromosomes is suppressed in hybrids by the *Xag* inversion, which is fixed within *An. coluzzii* but absent in *An. arabiensis.* The *Xag* inversion comprises ∼75% of the *An. coluzzii* X chromosome. This limits our ability to map genes to the X chromosome between species in the complex that harbor the *Xag* inversion (*An. gambiae, An. coluzzii,* and *An. merus*), and those that have the standard arrangement, including *An. quadriannulatus*.

Slotman et al. (2004) also identified both partially dominant and partially recessive male sterility QTL on the 2^nd^ and 3^rd^ chromosomes in both backcrosses. Consistent with the Dobzhansky-Muller model (Dobzhansky, 1936; Muller, 1940), and previous results from *Drosophila* (Wu and Beckenback, 1983; Orr and Coyne, 1989) the position of autosomal sterility QTL differs between each backcross. Complete sterility is achieved through the inheritance of at least three to four autosomal sterility loci (Slotman et al., 2004), including a pericentromeric QTL on the 3L chromosome arm. Hybrid male inviability is caused by an incompatibility between the *An. coluzzii* X chromosome and at least one locus on each autosome of *An. arabiensis*.

Female hybrid sterility QTL are present on the third chromosomes of both *An. coluzzii* and *An. arabiensis* (Slotman et al., 2005). A QTL on the second chromosome *An. coluzzii* causes female hybrid sterility when it is introgressed into an *An. arabiensis* genomic background. However, the female sterility phenotype is affected primarily by the interaction of a foreign X chromosome with the two autosomal QTL. Approximately 75% of observed phenotypic variance is due to the X chromosome, which only represents 8.8% of the genome (Holt et al., 2002). Due to the lack of recombination between X chromosomes in the cross, the number of loci on the X chromosome that effect the hybrid sterility phenotype is unknown. Therefore, the impact of the X on the hybrid sterility phenotype could result from one locus of major effect, or the effects could be spread across multiple loci. A comparison of the data between the male (Slotman et al., 2004) and female (Slotman et al., 2005) *An. coluzzii* × *An. arabiensis* hybrid sterility data shows that there are a greater number of male than female hybrid sterility QTL, supporting previous studies that found male sterility factors to evolve faster than female sterility factors (Charlesworth et al., 1987; Kirkpatrick and Hall, 2004). This is consistent with faster male evolution observed in *Aedes* mosquitoes (Presgraves and Orr, 1998), and the higher numbers of male versus female sterility factors in *Drosophila* (e.g. Hollocher et al., 1996; True et al., 1996; Tao and Hartl, 2003).

In order to better understand how post-zygotic reproductive isolation evolved during the rapid radiation of the of the *An. gambiae* species complex we performed a QTL analysis of male hybrid sterility in a (*An. coluzzii* × *An. quadriannulatus*) F1 × *An. quadriannulatus* backcross, and identified regions of the *An. coluzzii* genome that contribute to hybrid male sterility and inviability when introgressed into an *An. quadriannulatus* genetic background. We discuss how these QTL relate to those identified in the (*An. coluzzii × An. arabiensis*) F1 x *An. arabiensis* backcross (Slotman et al., 2004), and the implications these findings have on our understanding of the evolution of post-zygotic reproductive isolation in the *Anopheles gambiae* species complex, and more broadly, in the context of rapid species radiations.

## Methods

### Backcross and Phenotype Scoring

We performed a backcross between the SUA2La strain of *An. coluzzii* and the SANGUA strain of *An. quadriannulatus* to identify regions of the *An. coluzzii* genome that exhibit DMI with the *An. quadriannulatus* genome, resulting in F1 male sterility. The *An. coluzzii* strain was established from collections in Suakoko, Libera and the *An. quadriannulatus* strain was established in from populations in Sangwe, Zimbabwe. Both strains have been lab reared for hundreds of generations. We used a standard backcross scheme that has been used in studies to investigate the genetic basis of hybrid sterility and inviability going back to Dobzhansky (1936). F1 *An. coluzzii × An. quadriannulatus* (females × males, respectively) hybrids were reared and backcrossed to *An. quadriannulatus* males *en masse*. Consistent with crosses between *An. coluzzii* and *An. arabienesis*, the Xag inversion was expected to suppress recombination between the X chromosomes of *An. coluzzii* and *An. quadriannulatus*.

The *2La* inversion is also fixed between the mosquito strains used in this cross, limiting our mapping resolution in this region of the genome. While the *2La* is polymorphic within *An. coluzzii*, and *An. coluzzii* strains with the standard arrangement (co-linear with *An. quadriannulatus*) are available, we used the same *An. coluzzii* strain as Slotman et al. (2004) to allow a comparison between them.

The testes of male hybrids were dissected and squashed between a microscope slide and a cover slip. Each hybrid male mosquito was assigned a sterility phenotype following Slotman et al. (2004). Sterility scores were primarily based upon the proportion of normal to abnormal sperm. If no sperm was present, testes were often atrophied as well. In some instances, testes were missing completely. Phenotype scores were as follows: (1) normal testes and sperm, (2) slightly abnormal sperm, (3) 50/50 normal and abnormal sperm, (4) mostly abnormal sperm, (5) entirely abnormal sperm, (6) no sperm present and/or severely atrophied testes, (7) no testes present. Phenotyping was conducted by a single observer, without repeat observations. If unambiguous scoring was not possible, or if the reproductive tissues were damaged during dissection, the sample was discarded. Mosquitoes were preserved in 100% ethanol after phenotyping for DNA extraction an genotyping.

### DNA sequencing

DNA extractions were performed on a Qiagen Biosprint DNA extraction machine (Qiagen Inc, Valencia, CA). Extracted DNA was suspended in 200µl Qiagen elution buffer and stored at - 20°C. Each mosquito was genotyped using the restriction enzyme-based, genotype-by-sequencing approach according to the multiplexed shotgun genotyping (MSG) approach published by Andolfatto et al. (2011).

Briefly, 10 ng of genomic DNA of each mosquito was digested with the restriction enzyme MseI, which cuts the *An. gambiae* genome on average every 183.2 bp. Samples were digested for three hours at 37°C, followed by a 20 minute, 65°C enzyme deactivation step. Non-ligated bar-coded adapters were removed using an isopropanol precipitation, and bar-coded samples with unique adapters were pooled (96 per plate). Pooled samples were precipitated via centrifugation and re-suspended in TE buffer. Next, DNA was bead-purified using an Agencourt AMPure PCR purification kit. Bar-coded, purified DNA was size selected to a range of 250-400 bp using a Pippen Prep and then amplified using a Phusion High-Fidelity PCR kit. Samples were amplified using a common primer, and an indexed, library-specific primer (one per plate) that also incorporates the Illumina flow-cell adapter. Thus, individuals were identifiable to a library prep by their index and to the individual level by their barcoded adapter. Library amplification (PCR) was performed at a range of 12-18 cycles to identify the minimum number necessary to amplify the library to a target of 10-30 ng/uL (post clean up) in an effort to mitigate the incorporation of PCR errors. Amplified libraries were bead-purified using an Agencourt AMPure PCR purification kit. Library size distribution, integrity, and DNA concentration was confirmed by running each library on a NanoDrop and BioAnalyzer prior to sequencing. We phenotyped and sequenced a total of 841 male backcross hybrids. Hybrids were sequenced at the Texas A&M Genomics and Bioinformatics core facility on an Illumina HiSeq 2500 using 125 bp, single-end chemistry to 0.5-2.0x depth per individual.

We also performed pool-seq of ten *An. coluzzii* females and ten *An. quadriannulatus* females in order to identify fixed genetic difference between parental strains that would be diagnostic of the origin of genomic regions introgressed from *An. coluzzii* into *An. quadriannulatus* in the backcross. Libraries were sequenced on a single lane of Illumina HiSeq 2500 using 150 bp, paired-end chemistry to an average depth of 110.5x (*An. coluzzii*) and 66.9x (*An. quadriannulatus*). Additionally, we prepared and sequenced MSG libraries for 192 *An. coluzzii* and 192 *An. quadriannulatus* females to further validate ancestry informative markers and tune the MSG software error model.

### Parental pseudo-reference Construction

*An. coluzzii* pool-seq reads were aligned to the *An. gambiae* AgamP4.4 reference genome using the BWA mem alignment algorithm (Li and Durban, 2010) and SNPs were called using bcftools mpileup (v1.3) (Li et al., 2009). SNPs within three bases of indels were excluded. A pseudo-reference for *An. coluzzii* was created by updating the AgamP4.4 reference genome (vectorbase.org) with SNPs that had read depth between greater than 10 and less than 1000, map quality greater than 20, and a phred-scaled quality score for the alternative allele greater than 26. The *An. quadriannulatus* pseudo-reference was created in a similar fashion, but by aligning it to the *An. coluzzii* pseudo-reference and updating this with fixed SNPs (*An. coluzzii* – *An. quadriannulatus* divergent) that met the thresholds above.

### Hybrid Genotyping

We used Trimmomatic version 0.30 (Bolger et al., 2014) to trim MSG sequencing reads by quality by simultaneously soft clipping the reads from both 5’ and 3’ ends to an average phred quality score of 20, with no single bp in a four bp window below a phred quality score of 20. Only reads ≥ 50 bp were retained for alignment and subsequent analyses.

We used the MSG software package (Andolfatto et al., 2011) to align hybrid MSG data to parental pseudo-references and estimate the posterior probabilities of ancestry at genome-wide SNPs that were informative of parental ancestry. We removed 33 hybrids from this analysis due to low sequencing coverage (<10K reads). Posterior probabilities of ancestry were thinned using pull_thin_tsv.py (https://github.com/dstern/pull_thin). We considered a marker to be informative if its conditional probability of being homozygous differed by more than 0.10 from its neighboring marker. Missing genotypes were imputed using the hidden Markov model implemented in the sim.geno() function of R/qtl (Broman et al., 2003)

### Sterility QTL Mapping

We used R/qtl (Broman et al., 2003) to identify redundant, adjacent markers and further thinned our marker set to include only informative markers. It is important to note that this experiment is not marker, but recombination limited. Therefore, many markers are redundant, and we removed these to reduce computational time. Next, we used R/qtl to estimate the genetic map of 808 hybrid males using the Kosambi map function (Kosambi, 1943). We performed quantitative trait locus mapping with the composite interval mapping (CIM) method using 15 markers as co-factors, and also assessed the significance of QTL models using multiple imputation mapping. This method drops one QTL at a time to calculate the probability of QTL locations and their interactions using an ANOVA (Sen and Churchill, 2001). In all analyses, sterility was treated as a normal trait, and we used 1000 permutation replicates to assess the significance of log of odds (LOD) scores.

Ultimately, we took three approaches to identify sterility QTL in the dataset. The first was to scan for QTL using the composite interval mapping and calculating the logarithm of these odds (LOD) score every 1 cM. When calculating LOD scores we controlled for the genetic background of adjacent markers by using 15 marker covariates across the genome. Due to the skew of the phenotype scores toward “normal”, and severe segregation distortion of the X chromosome, we hypothesized that the QTL identified through CIM of the full dataset were in fact comprised of two types: (1) autosomal *An. coluzzii* loci that are incompatible with the *An. quadriannulatus* X, and (2) homozygous *An. quadriannulatus* autosomal loci that interact epistatically with the *An. coluzzii* X in individuals that are otherwise primarily heterozygous at autosomal loci (recovering viability despite the inheritance of a non-recombinant *An. coluzzii* X chromosome). Thus, we also split the dataset into individuals that had inherited the *An. quadriannulatus* X (N = 568) and *An. coluzzii* X (N = 211), respectively, and performed CIM (as above) on these datasets separately.

Next, we tested all peaks identified by CIM, regardless of significance or LOD score, in the full dataset, and any peaks that were unique to either the *An. quadriannulatus* X or *An. coluzzii* X datasets, respectively, by adding each peak independently to the multiple imputation QTL model that also contained the highly significant X chromosome QTL. After identifying all peaks that had a significant additive effect in the multiple imputation QTL model, we tested epistatic interactions between each, in a pairwise fashion, by adding each of these interactions to the model independently. This approach allowed us to identify peaks (and their interactions) which, despite having non-significant LOD scores in the CIM permutation test, were found to have a significant impact on sterility when incorporated into the multiple imputation QTL model.

We identified candidate sterility genes within QTL by determining the 1.5 LOD interval of all QTL peaks in base pairs, and used these coordinates to download the list of annotated genes within QTL from the AgamP4 genome using the VectorBase (vectorbase.org) genome browser using the “Export Data” function. Next we used the PANTHER classification system (Mi et al., 2019) gene list analysis search function (pantherdb.org) to search for the protein annotations and functions of these genes. Finally, we performed a protein network analysis on the entire gene list using STRING (Szklarczyket al., 2019) in order to identify direct interactions between putative sterility genes.

### Identification of Inviability Loci

We identified regions of the *An. coluzzii* genome that result in hybrid inviability when introgressed into an *An. quadriannulatus* genetic background by analyzing segregation distortion of autosomal loci in the *An. quadriannulatus* X chromosome and *An. coluzzii* X chromosome datasets. We used the mean, genome-wide proportion of *An. quadriannulatus* homozygosity in the full *An. coluzzii × An. quadriannulatus* backcross population as the expected homozygosity proportion, assuming Hardy-Weinberg equilibrium. We used a chi-square test to identify loci that deviate significantly (p-value < 6.01e-05 after Bonferroni correction) from expected homozygosity.

## Results

### Pseudo-reference Construction, Genotyping, and Genetic Map

After whole genome alignment, SNP calling, and variant filtering of the *An. coluzzii* pool-seq data, we generated the *An. coluzzii* pseuo-reference by incorporating 4,391,200 filtered SNPs into the *An. gambiae s.s.* AgamP4 reference genome. To construct the *An. quadriannulatus* pseuo-reference, we updated the *An. coluzzii* pseudo-reference with 4,242,426 *An. quadriannulatus* SNPs that were found to be fixed between the *An. quadriannulatus* and *An. coluzzii* pool-seq samples. MSG data is inherently low coverage, so not all fixed differences between the pool-seq datasets were subsequently used for hybrid genotyping.

We calculated genotype probabilities at 239,420 ancestry informative SNPs for 808 *An. coluzzii × An. quadriannulatus* backcross hybrid males. Ultimately, we trimmed the dataset to 860 non-redundant markers (Table 1). The mean proportion of missing data per individual was 0.021 (sd = 0.060, range = 0.0 – 0.445). These markers were used to calculate a genetic map (Figure 1) with Kosambi distances of 206.1 cM for the second chromosome (110.9 Mb), 100.7 cM for the third (95.2 Mb), and 3.7 cM for the X (24.4 Mb) (Table 1, Figure 1). The larger relative genetic map of the second chromosome is likely due to the presence of the *2La* inversion, which is fixed between these species and can significantly increase recombination frequency on the 2R (i.e. Schultz-Redfield Effect, Schultz and Redfield, 1951; Lucchesi and Suzuki, 1968; Stevison et al., 2011), and segregation distortion resulting from male inviability for some genotype combinations (Zuo et al., 2019; Hackett and Broadfoot, 2003).

**Figure 1.**
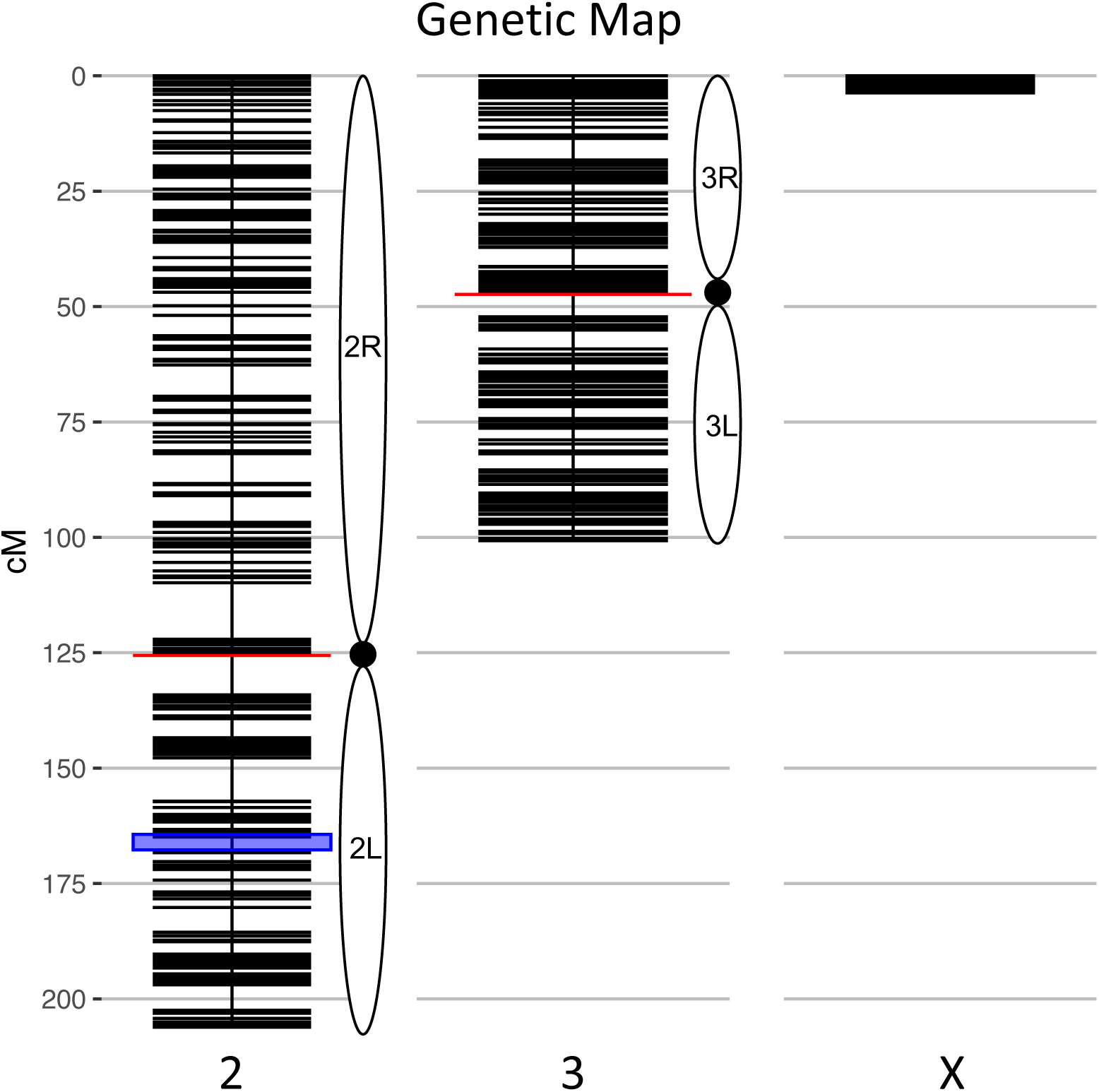
Genetic map of the *An. coluzzii × An. quadriannulatus* backcross showing centromere locations (red horizontal lines) and marker locations (black horizontal lines). The centromere of the telocentric X chromosome is opposite zero cM. The blue highlighted area on chromosome 2 indicates the location of the 2L*a* inversion in *An. coluzzii*.

**Table 1.** The number of ancestry informative, thinned, and non-duplicate markers genotyped in hybrids, and the physical and map distances of each chromosome in the cross.

| **Chromosome** | **Informative**<br>**Markers** | **Thinned**<br>**Markers** | **Non-duplicate**<br>**markers** | **Physical Length**<br>**(Mb)** | **Genetic Length**<br>**(cM)** |
| --- | --- | --- | --- | --- | --- |
| 2 | 120,717 | 710 | 440 | 110.9 | 206.1 |
| 3 | 97,706 | 629 | 392 | 95.2 | 100.7 |
| X | 20,997 | 217 | 28 | 24.4 | 3.7 |

The genetic length of the X chromosome was expected to be low due to the presence of the *Xag* inversion, which comprises 80% of the X and is fixed for alternate arrangements between these species. The majority of hybrids (70.3%, N = 568) inherited a non-recombinant *An. quadriannulatus* X chromosome, 26.1% (N = 211) inherited a non-recombinant *An. coluzzii* X, and 3.5% (N = 29) a recombinant X. The mean sterility score (ranging from 1-7, normal to completely sterile) was 2.62 (sd = 2.34). The majority of backcross mosquitoes (64.23%, N =519) were fully fertile (sterility score = 1).

### Segregation distortion and inviability

Segregation distortion was previously documented in male and female hybrids between member species of this complex (Slotman et al., 2004; Slotman et al., 2005; Smith et al., 2015). Furthermore, the complete absence of specific genotypes can indicate their inviability. We tested for significant segregation distortion of autosomal loci in the *An. coluzzii* X and *An. quadriannulatus* X datasets, respectively, be comparing the proportion of *An. quadriannulatus* homozygosity at each locus to the genome-wide average in the full dataset. It is important to note that this analysis was performed on a per locus basis, and is agnostic of interactions between loci.

In the *An. quadriannulatus* X dataset, 67.3% (N=296) of chromosome 2 loci and 18.4% (N=72) of chromosome 3 loci showed significant segregation distortion (Figure 2). Loci on the chromosome 2 exhibit both homozygote excess and deficit (heterosis), while only homozygote excess is exhibited on chromosome 3. In the *An. coluzzii* X dataset, 23.9% (N=105) of chromosome 2 loci and 69.6% (N=273) of chromosome 3 loci showed significant segregation distortion. Interestingly, 40.8% of chromosome 3 loci (spanning a 18.9 Mb region) lack homozygotes in all genotyped individuals, indicating individuals that inherited the *An. coluzzii* X and were *An. quadriannulatus* homozygous at loci inside this region were completely inviable (Figure 2). No autosomal heterozygote excess was observed in the *An. coluzzii* X dataset.

**Figure 2.**
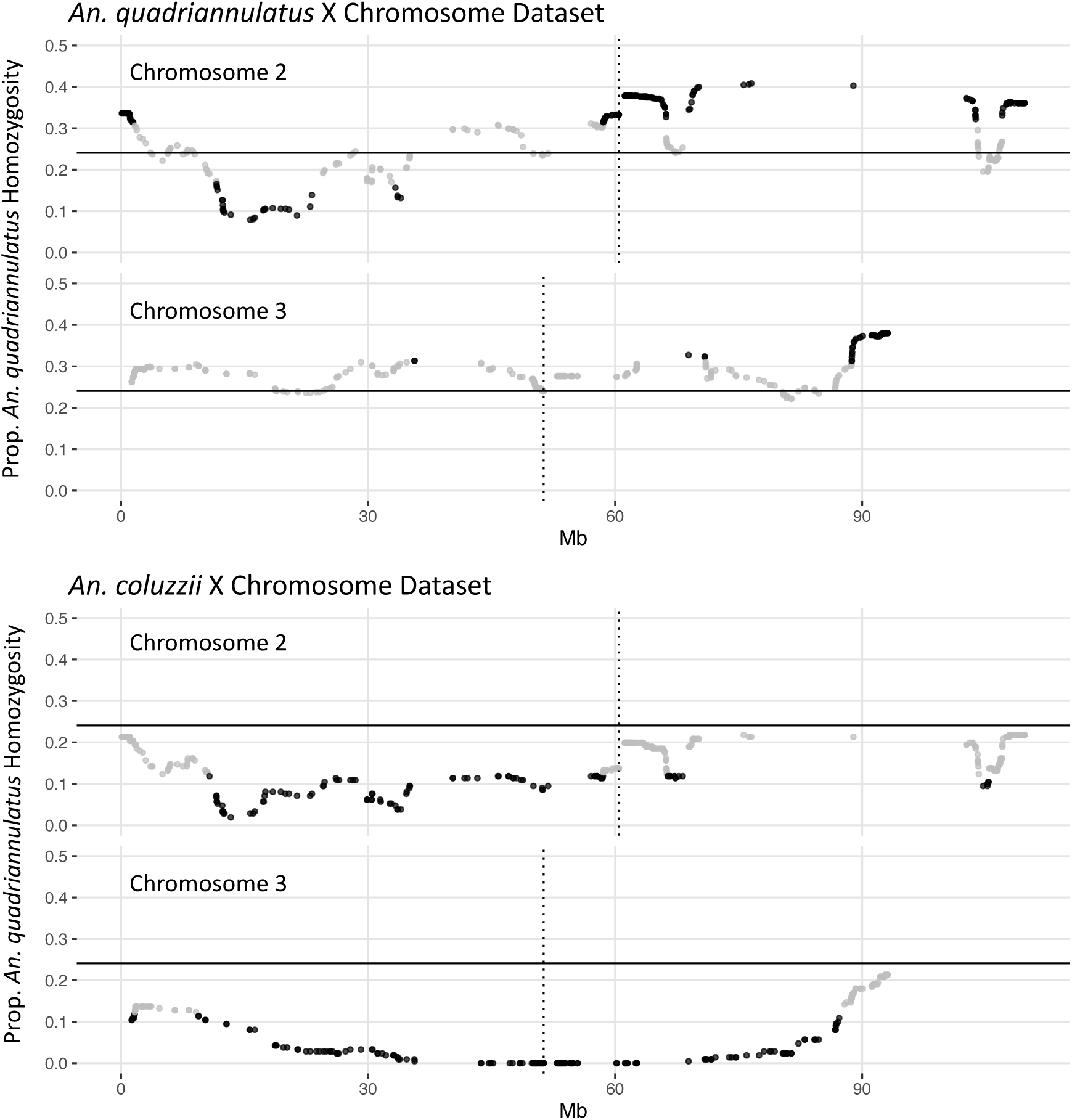
Segregation distortion at chromosome 2 and 3 loci for the *An. quadriannulatus* X chromosome (top) and *An. coluzii* X chromosome dataset (bottom). Grey dots indicate SNPs that do not deviate significantly from the expected proportion of homozygosity (solid horizontal line), while black dots deviate significantly (chi-square p-val < 0.05). Vertical dotted lines indicate centromere locations.

### Composite Interval Mapping

Due to minimal recombination events, the X chromosome is effectively one large QTL with a LOD score of 203.4 (Table 2). By comparison, none of the autosomal QTL reach a LOD score of 4.0. No autosomal QTL fall above the CIM 5% significance threshold (LOD=4.31, 1,000 permutations). Despite this, we annotated three QTL peaks on the second chromosome (1.63 cM, 20.87 cM, and 134.26 cM) (Figure 3, Table 2) that contributed significantly to the sterility phenotype when included in the multiple imputation QTL model (see *Multiple Imputation*, below, and Table 4). The second chromosome peak at 1.63 cM (QTL “2R-A”) was present in the full and *An. quadriannulatus* X datasets, but not the *An. coluzzii* X dataset (Figure 3). QTL “2L” at 134.26 cM was present in the full and *An. coluzzii* X datasets, but not the *An. quadriannulatus* X dataset. Finally, QTL “2R-B” (20.87 cM) was found in the full and both X chromosome split datasets, suggesting that it contains loci involved in autosomal-autosomal incompatibilities (Figure 3). While additional LOD peaks are present in the *An. coluzzii* X dataset, these were non-significant when included in the multiple imputation QTL model (see *Multiple Imputation*, below) (Figure 3). None of the second chromosome QTL overlap with those found in the *An. coluzzii × An. arabiensis* cross (Slotman et al., 2004) (Figure 3, Table 2).

**Figure 3.**
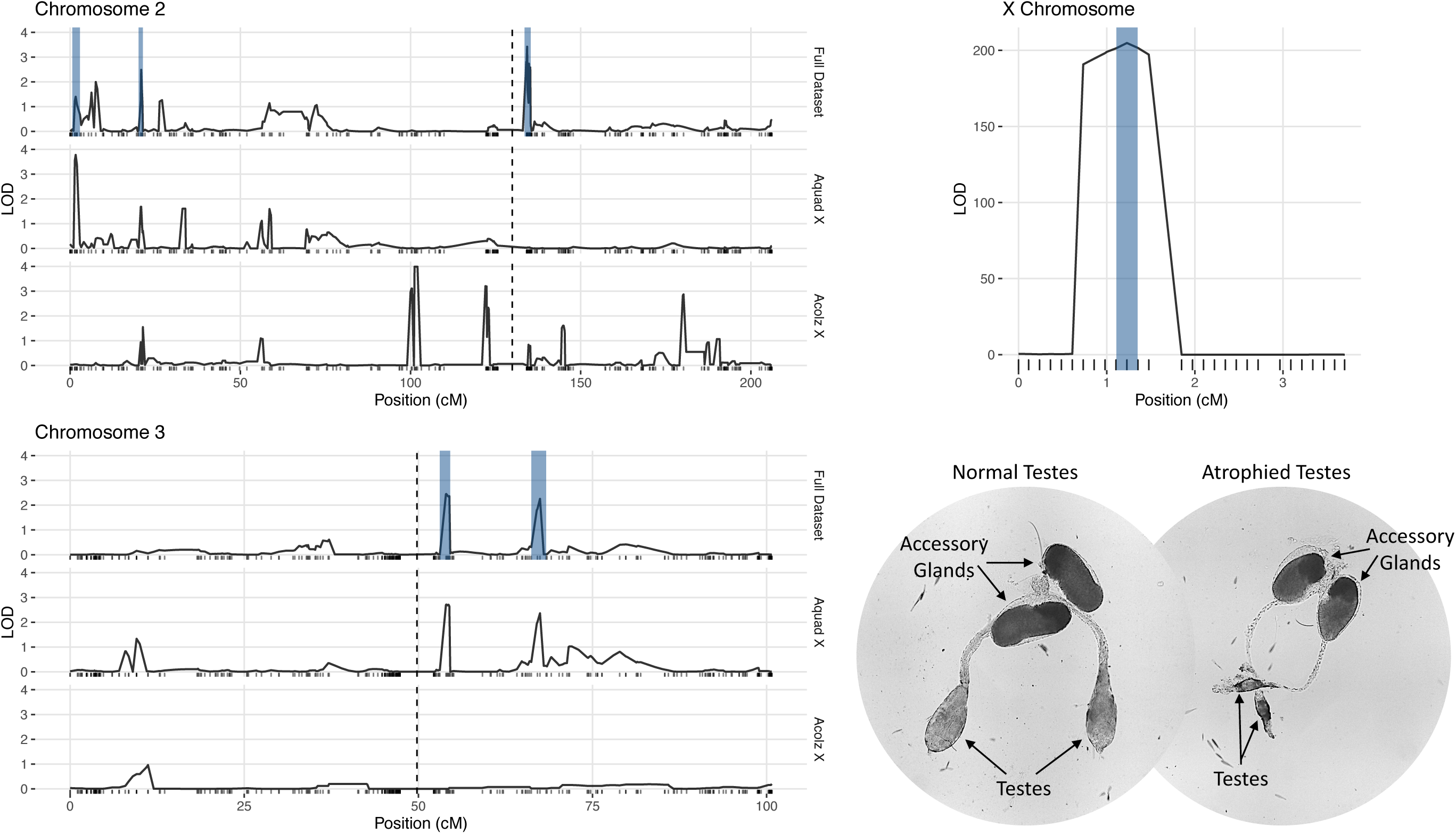
LOD plots for hybrid male sterility QTL identified with composite interval mapping in the *An. coluzzii × An. quadriannulatus* cross. Note the differences in the y-axis scales between the chromosome 2 (top left) and 3 (bottom left) panels, and the X chromosome panel (top right). Vertical, dashed lines indicate centromere locations on chromosomes 2 and 3. Second and third chromosome right arms are left of the centromere, and left arms are to the right. Vertical blue bars indicate 1.5 LOD intervals of QTL that explain a significant proportion of phenotypic variation in multiple imputation QTL model of the full dataset. The bottom right panel illustrates the gross morphological differences between hybrid testes with a normal sterility score (1) and a atrophied hybrid testes (sterility score = 6) which contain no sperm or only a few sperm with abnormal morphology.

**Table 2.** Genomic regions of QTL in physical (Mb) and map (cM) distances. Start an end positions indicate the 1.5 LOD intervals of QTL locations (LOD peak). Bold values indicate overlapping QTL between the crosses on the 3L, a region of the *An. coluzzii* genome that contributes to hybrid male sterility when introgressed into both *An. quadriannulatus* and *An. arabiensis* genetic backgrounds.

| Cross | QTL | LOD | LOD peak (cM) | Start (cM) | End (cM) | Length (cM) | LOD peak (Mb) | Start (Mb) | End (Mb) | Length (Mb) | Num Genes |
| --- | --- | --- | --- | --- | --- | --- | --- | --- | --- | --- | --- |
| *An. coluzzii* ×<br>*An. quadriannulatus* | X | 204.84 | 1.23 | 1.11 | 1.35 | 0.25 | 6.68 | 2.96 | 11.48 | 8.53 | 453 |
|  | 2R-A | 3.78 | 1.63 | 0.63 | 2.88 | 2.26 | 1.15 | 1.03 | 1.67 | 0.63 | 49 |
|  | 2R-B | 2.49 | 20.87 | 20.12 | 21.37 | 1.24 | 8.11 | 8.03 | 8.73 | 0.70 | 17 |
|  | 2L | 3.42 | 134.26 | 134.13 | 135.30 | 1.18 | 2.69 | 0.73 | 4.14 | 3.42 | 153 |
|  | 3L-A | 2.45 | 53.95 | 53.08 | 54.57 | 1.49 | 11.18 | **10.18** | **11.30** | 1.13 | 52 |
|  | 3L-B | 2.26 | 67.46 | 66.21 | 68.34 | 2.12 | 22.82 | 20.86 | 23.33 | 2.47 | 78 |
| *An. coluzzii* ×<br>*An. arabiensis*<br>(Slotman *et al.*, 2004) | 2R | 38.30 | 16.07 | 15.25 | 18.05 | 2.80 | 40.44 | 33.18 | 48.30 | 14.83 | - |
|  | 3R | 3.69 | 12.82 | 8.41 | 19.04 | 10.63 | 14.83 | 8.79 | 44.24 | 23.58 | - |
|  | 3L | 7.03 | 23.23 | 19.04 | 26.08 | 7.04 | 16.86 | **44.24 (3R)** | **19.77** | 38.37 | - |

Two QTL were identified on the third chromosome at 53.95 (QTL “3L-A”) and 67.46 cM (QTL “3L-B”) (Figure 3, Table 2). These are both present in the full and *An. quadriannulatus* X dataset, but not the *An. coluzzii* X dataset, indicating that *An. coluzzii* introgressions into a predominantly *An. quadriannulatus* genetic background are driving this signal. QTL “3L-A” in the *An. coluzzii × An. quadriannulatus* dataset overlaps with a QTL in the *An. coluzzii × An. arabiensis* dataset (Slotman *et al*, 2004) (Figure 3, Table 2). The 3L-A and 3L-B QTL have seemingly additive-only effects on sterility (see *Multiple Imputation*, below); these QTL were not identified by CIM in the *An. coluzzii* X dataset, suggesting that they result only from *An. coluzzii* introgressions into predominantly *An. quadriannulatus* genetic backgrounds. However, only three hybrid males were sampled that are *An. quadriannulatus* 3L-B homozygous and inherited the *An. coluzzii* X (Figure 4), and each had a sterility score of five or six. No hybrid males were sampled that are *An. quadriannulatus* 3L-A homozygous and inherited the *An. coluzzii* X. The reduction (or absence) of homozygous genotypes at the 3L-A and 3L-B QTL differs significantly from expectation (the percentage of homozygous genotypes across all autosomal loci, assuming HWE) (Table 3). Thus, these regions of the *An. quadriannulatus* genome interact epistatically with the X and result in almost complete male inviability when introgresesed into an *An. coluzzii* genetic background.

**Figure 4.**
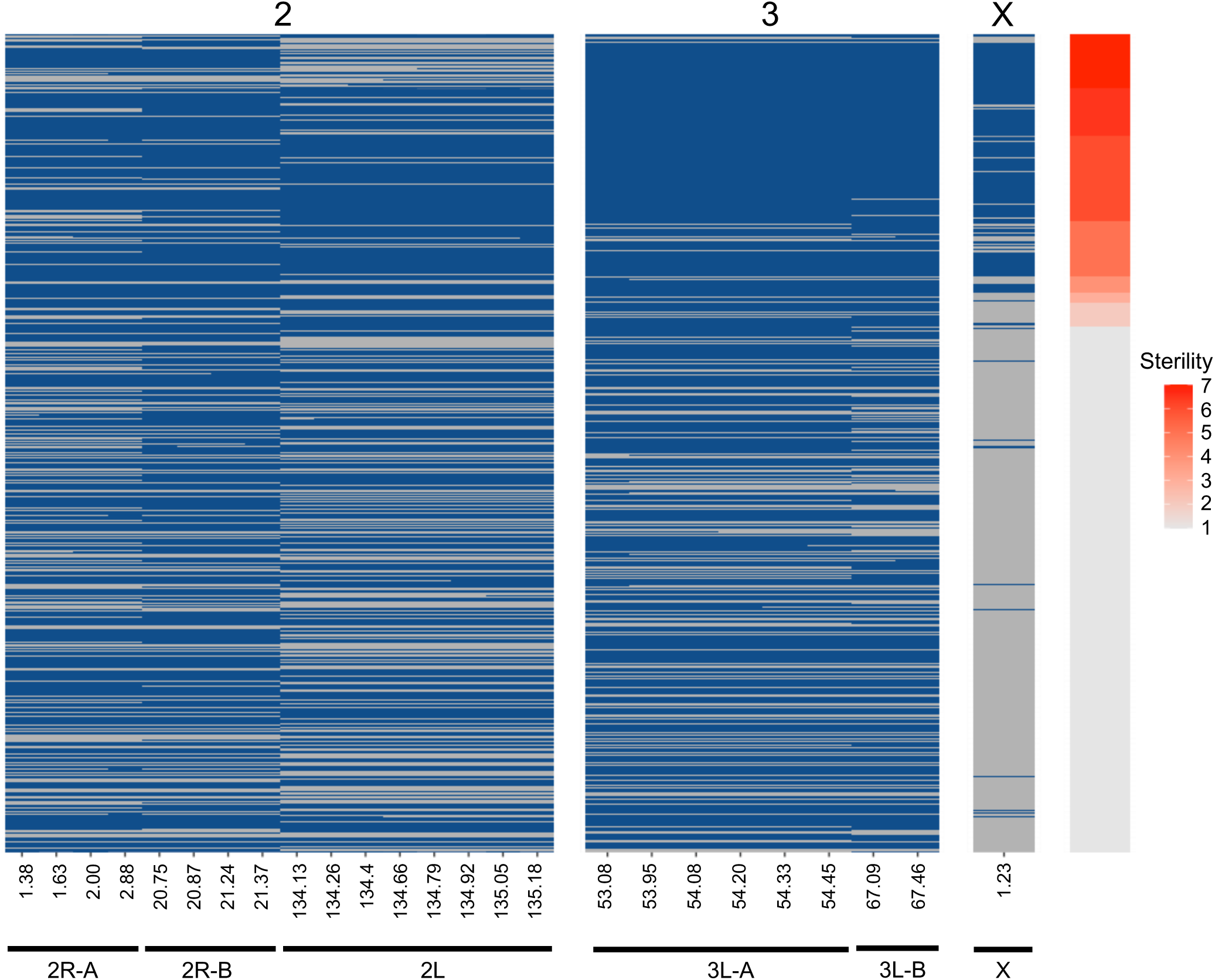
Genotype heat map for each *An. coluzzii × An. quadriannulatus* backcross male hybrid (y-axis) at the six QTL (x-axis) identified using CIM and multiple imputation. All markers within the 1.5 LOD interval of each QTL peak (Table 2) are included (x-axis, positions in cM). Individuals are sorted on the y-axis by sterility, with the individuals with the highest sterility score (7) at the top. Blue cells indicate *Aquad/Acolz* heterozygosity on the autosomes, and *Acolz* hemizygosity on the X chromosome. Grey cells indicate *Aquad/Aquad* homozygosity on the autosomes, and *Aquad* hemizygosity on the X chromosome.

**Table 3.** Percentage of *An. quadriannulatus* homozygosity at hybrid male sterility autosomal QTL peaks in the full, *An. quadriannulatus* X chromosome, and *An. coluzzii* X chromosome datasets. p-values are reported for chi-square tests, which were used to test for significant deviation (p-value < 6.01e-05 after Bonferroni correction) from expected homozygosity. Expected homozygosity was calculated as the mean homozygosity percentage across all autosomal loci in the full dataset, assuming Hardy-Weinberg equilibrium (0.2409).

| QTL | cM | Full dataset |  | *An. quadriannulatus* X |  | *An. coluzzii* X |  |
| --- | --- | --- | --- | --- | --- | --- | --- |
| | | Homozygosity (%) | $\chi^2$ p-val | Homozygosity (%) | $\chi^2$ p-val | Homozygosity (%) | $\chi^2$ p-val |
| 2R-A | 1.63 | 28.5 | 3.7E-03 | 32.0 | 9.5E-06 | 20.4 | 2.1E-01 |
| 2R-B | 20.87 | 22.2 | 2.0E-01 | 24.8 | 6.8E-01 | 16.1 | 6.7E-03 |
| 2L | 134.26 | 32.1 | 1.2E-07 | 37.7 | 3.8E-14 | 19.9 | 1.5E-01 |
| 3L-A | 53.95 | 20.4 | 1.5E-02 | 29.0 | 5.8E-03 | 0.0 | 2.7E-16 |
| 3L-B | 67.46 | 20.9 | 3.5E-02 | 29.0 | 5.8E-03 | 1.4 | 1.3E-14 |

### Multiple Imputation

CIM detects QTL by calculating LOD scores at each marker or map location along each chromosome and does this agnostic of other QTL contributions to the phenotype and potential epistatic interactions between QTL. Due to this limitation, we built multiple imputation QTL models to detect the significance of all QTL LOD peaks found during CIM of the full (i.e., not split by X chromosome) dataset. The QTL discussed above, and those reported in Table 2, are only those that contributed significantly to the variance in sterility in the multiple imputation models described herein. We report two QTL models, one that considers only additive effects of QTL on sterility, and one which includes significant epistatic effects between QTL. We took an iterative process during testing QTL models and added QTL locations identified through CIM in a stepwise fashion. After we had identified those that had significant additive effects, we tested all pairwise epistatic effects between these loci. The final “additive only” and “epistasis allowed” models explain 71.66% and 72.29% of the variance in sterility, respectively (Table 4). The vast majority of this variance is explained by the inheritance of the X chromosome. As expected, individuals harboring the *An. coluzzii* X chromosome have on average higher sterility scores because in these backcrosses there are more opportunities for *An. quadriannulatus* autosomal – *An. coluzzii* X incompatibilities than vice versa (Figure 4).

**Table 4.**
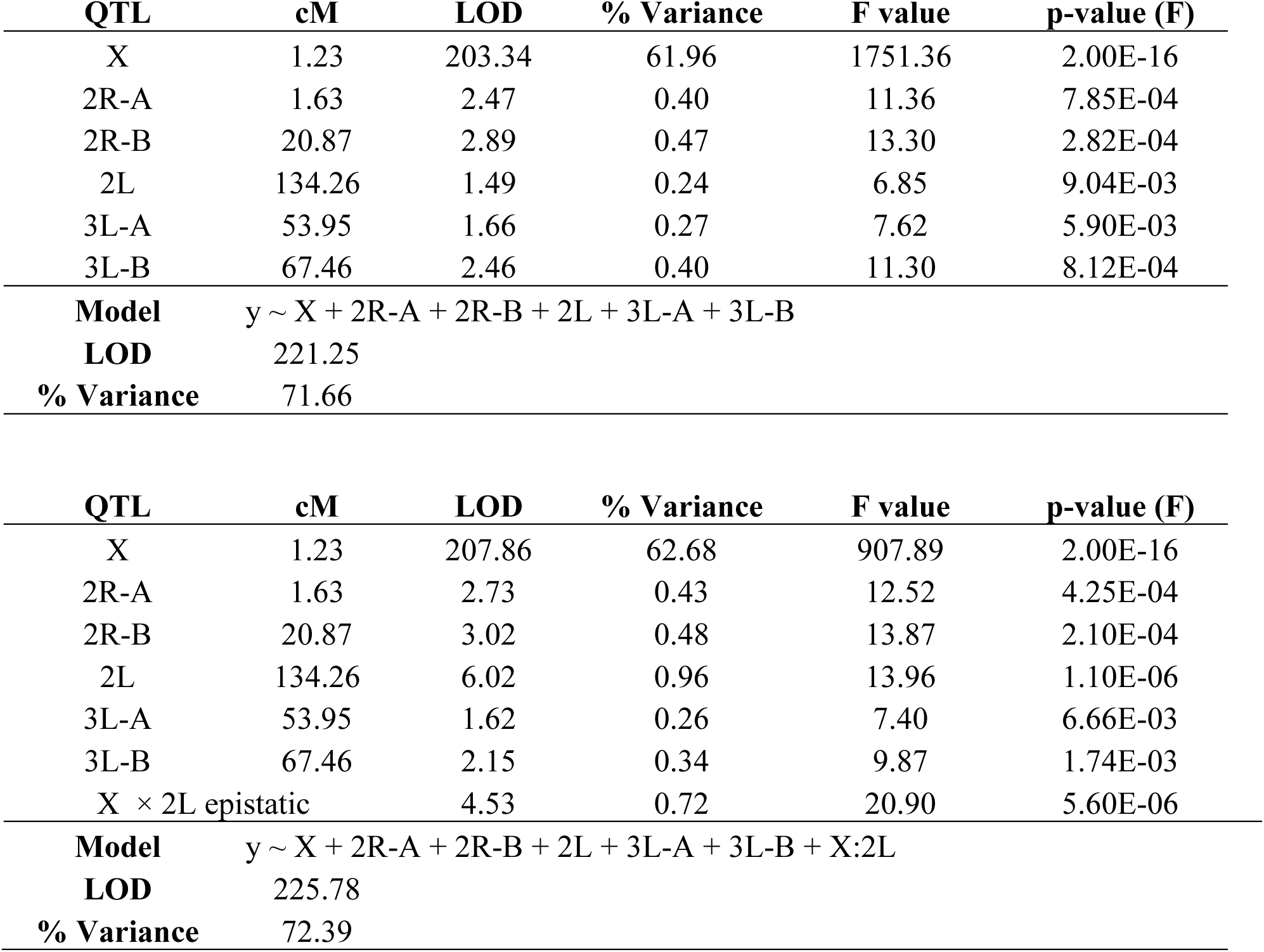
Results of two multiple imputation QTL models (ANOVA). The top model includes only additive model includes all additive effects and also an epistatic effect between the X chromosome and the 2L QTL.

Each of the QTL described had a significant additive effect in both the “additive only” and “epistasis allowed” models (F-test p-value < 0.01) (Table 4). After testing all possible pair-wise interactions between these loci, we found that only the X and 2L (134.26 cM) QTL pair had a significant interaction (F-test p-value = 5.60E-06). In both models, additive effects of autosomal loci, and the epistatic interaction between the X and 2L QTL, each explained less than 1% of the variance in hybrid sterility. The “epistasis allowed” model explained a higher proportion of the variance and had a higher model LOD (225.78) when compared to the “additive only” model (221.25) (Table 4).

To better understand the direction of the effect of each QTL in the full dataset, we compared the phenotypes of each individual given their genotype (*An. quadriannulatus* homozygous (QQ) vs. *An. quadriannulatus/An. coluzzii* heterozygous (QC)) at autosomal QTL and the X chromosome (Figure 5). Each autosomal locus increases sterility in the heterozygous state, and individuals with the *An. coluzzii* X are significantly more sterile. The interaction plot (Figure 5, “X vs 2L”, right panel) between the X and 2L QTL demonstrates that heterozygosity at the 2L QTL alone does not impact sterility when an individual has the *An. quadriannulatus* X, but reduces sterility when inherited with the *An. coluzzii* X. The evidence suggests that each autosomal QTL indicate the location of a (partially) dominant *An. coluzzii* sterility gene that causes incompatibilities with the *An. quadriannulatus* background.

**Figure 5.**
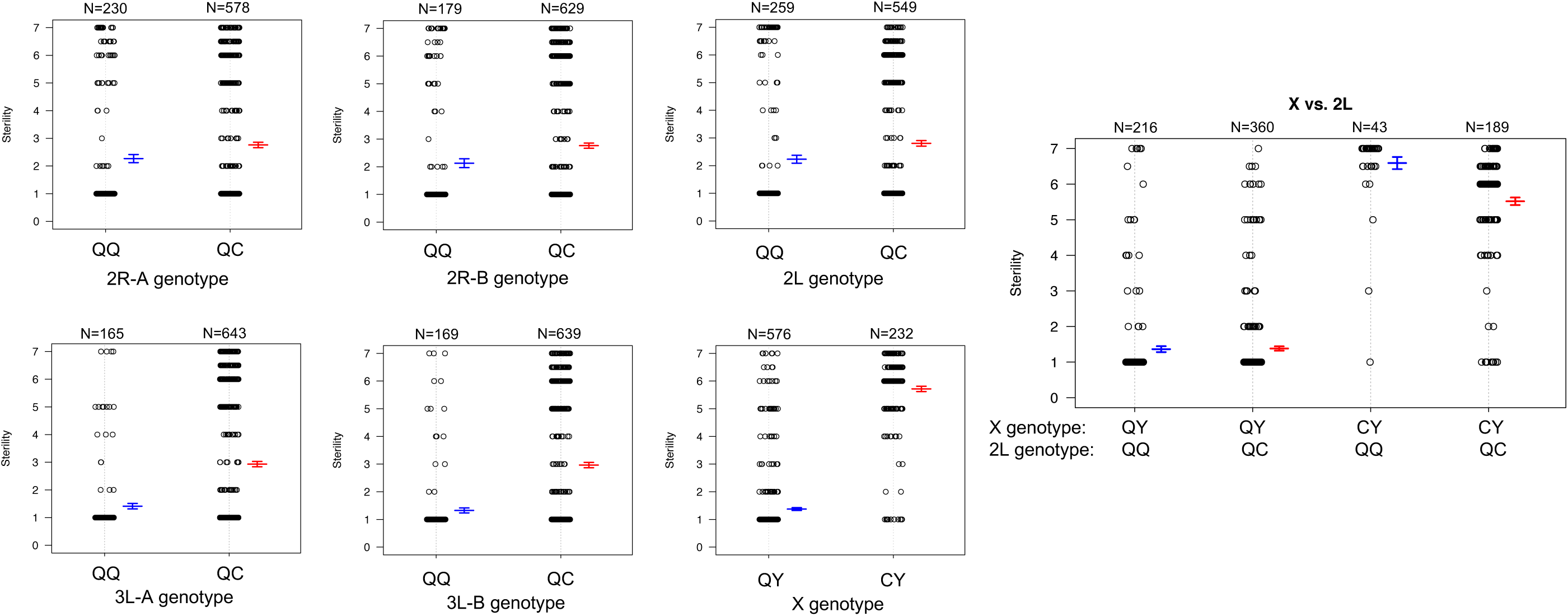
Genotype (x-axis) by phenotype (sterility score, y-axis) plots for the six QTL identified using CIM and multiple imputation, and an interaction plot (right) for the epistatic effect between the X chromosome and 2L QTL. Each individual is represented as a dot, which are jittered to avoid overlap. Box plots illustrate the mean (middle line) of each phenotype distribution and whiskers illustrate +/-1 standard error of the mean. QTL names and locations correspond to those in Table 2. QQ: *An. quadriannulatus* homozygous, QC: *An quadriannulatus / An. coluzzii* heterozygous.

We determined the 1.5 LOD interval of each QTL and, using the base pair positions of our SNP markers, determined the physical map span of each (Table 2). Next, we compared QTL locations to male sterility QTL found in a *An. coluzzii* × *An. arabiensis* cross (Figure 3) (Slotman et al., 2004). We re-analyzed this data using CIM with three marker covariates. No significant QTL on the second chromosome overlap between these crosses. However, the 3L-A QTL in our *An. coluzzii × An. quadriannulatus* cross falls within the 3L QTL from the *An. coluzzii × An. arabiensis* cross, and the 3L-B QTL overlaps it. In the Slotman et al. (2004) data the third chromosome QTLs are also only present when the hybrid has the backcross (*An. arabiensis* in this case) parent X chromosome (Figure 3). Thus, interestingly this represents a region of the *An. coluzzii* that contributes to sterility when introgressed into both *An. quadriannulatus* and *An. arabiensis* genetic backgrounds.

### Candidate Speciation Genes

We identified all annotated genes in the AgamP4 genome within the 1.5 LOD intervals of each QTL using VectorBase (www.vectorbase.org) and found the protein family (if available) of each gene using PANTHER. Due to the lack of resolution on the X chromosome, we excluded this chromosome from this analysis. We identified 349 genes within all autosomal QTL (2.47 Mb total, Table 2). We identified ten genes with protein functions related to mating, reproduction, testes, and microtubule morphogenesis and function (Table 5) in the full gene set (Table S1).

**Table 5.** Annotated genes in the AgamP4 genome within autosomal QTL with functions related to mating, reproduction, and sperm motility and morphogenesis. Gene names in bold indicate those interacting in a regulatory network associated with sperm microtubule morphogenesis and function. STRING network analysis output can be found in Supplementary File 1: Deitz_et_al_2020_String_protein_interaction_network.xlsx.

| QTL | Gene Name | Panther Gene Ontology<br>Family / Subfamily | Protein Description | Protein Description<br>Source |
| --- | --- | --- | --- | --- |
| 2R-A | **AGAP001229** | DYNEIN LIGHT CHAIN<br>TCTEX-TYPE<br>(PTHR21255:SF4) | Tctex-1 (t-complex testis expressed-1) is a dynein light chain. Dynein translocates rhodopsin-bearing vesicles along microtubules. In mice, the chromosomal location and pattern of expression of Tctex-1 make it a candidate for involvement in male sterility. | https://www.ebi.ac.uk/interpro/entry/InterPro/IPR005334/ |
| 2R-A | AGAP001227 | GAMMA-TUBULIN<br>COMPLEX<br>COMPONENT 5<br>(PTHR19302:SF33) | The microtubule organizing centers (MTOCs) of eukaryotic cells are the sites of nucleation of microtubules, and are known as the centrosome in animal cells and the spindle pole body in yeast. Gamma-tubulin, which is 30% identical to alpha and beta tubulins that form microtubules, appears to be a key protein involved in nucleation of microtubules. | https://www.ebi.ac.uk/interpro/entry/InterPro/IPR007259/ |
| 2R-A | AGAP001194 | MICROTUBULE-ASSOCIATED PROTEIN<br>FUTSCH<br>(PTHR13843:SF12) | futsch (futsch) encodes a microtubule binding protein involved in the formation of synaptic buttons at the neuromuscular junctions. Its expression levels are regulated by the product of TBPH to prevent synaptic defects. | https://flybase.org/reports/FBgn0259108 |
| 2R-A | **AGAP001219** | TUBULIN-LIKE<br>PROTEIN ALPHA-4B-RELATED<br>(PTHR11588:SF239) | Microtubules are polymers of tubulin, a dimer of two 55kDa subunits, designated alpha and beta. Within the microtubule lattice, alpha-beta heterodimers associate in a head-to-tail fashion, giving rise to microtubule polarity. Fluorescent labelling studies have suggested that tubulin is oriented in microtubules with beta-tubulin toward the plus end. | https://www.ebi.ac.uk/interpro/entry/InterPro/IPR002452/ |
| 2L | AGAP004767 | TRANSCRIPTION<br>FACTOR KEN<br>(PTHR45993:SF7) | Mutations in the ken and barbie locus are accompanied by the malformation of terminalia in adult *Drosophila*. Male and female genitalia often remain inside the body, and the same portions of genitalia and analia are missing in a fraction of homozygous flies. Rotated and/or duplicated terminalia are also observed. Terminalia phenotypes are enhanced by mutations in the gap gene *tailless*, the homeobox gene *caudal*, and the decapentaplegic gene that encodes a TGFbeta-like morphogen. The ken and barbie gene encodes a protein with three CCHH-type zinc finger motifs that are conserved in several transcription factors such as Krüppel and BCL-6. | https://flybase.org/reports/FBgn0011236 |
| 2L | **AGAP004724** | INTRAFLAGELLAR<br>TRANSPORT PROTEIN<br>74 HOMOLOG<br>(PTHR31432:SF0) | Intraflagellar transport of ciliary precursors such as tubulin from the cytoplasm to the ciliary tip is required for ciliogenesis. Intraflagellar transport protein 74 (IFT74) is a component of the intraflagellar transport (IFT) complex B, which contains at least 20 different protein subunits. Together with IFT81, it forms a tubulin-binding module that specifically mediates transport of tubulin within the cilium. | https://www.ebi.ac.uk/interpro/entry/InterPro/IPR029602/ |
| 2L | AGAP004817 | PROTEIN LINGERER<br>(PTHR16308:SF13) | lingerer (lig) encodes a putative RNA binding protein that forms a complex with the products of orb and orb2. Loss of lig results in defects in copulation and short-term memory in *Drosophila*. | https://flybase.org/reports/FBgn0020279 |
| 2L | **AGAP004782** | TUBULIN-SPECIFIC<br>CHAPERONE D<br>(PTHR12658:SF0) | The tubulin heterodimer consists of one alpha- and one beta-tubulin polypeptide. Tubulin-specific chaperones are essential for bring the alpha- and beta-tubulin subunits together into a tightly associated heterodimer. | https://www.ebi.ac.uk/interpro/entry/InterPro/IPR033162/ |
| 3L-A | **AGAP010790** | CYTOPLASMIC<br>DYNEIN 2 HEAVY<br>CHAIN 1<br>(PTHR10676:SF352) | Cytoplasmic dynein 2 heavy chain 1 (DYNC2H1) is a subunit of the cytoplasmic dynein complex 2. It may function as a motor for intraflagellar retrograde transport. It also plays a role in cilia biogenesis. | https://www.ebi.ac.uk/interpro/entry/InterPro/IPR026815/ |
| 3L-B | AGAP011361 | CHECKPOINT PROTEIN<br>(PTHR12900:SF0) | Hus1-like (Hus1-like) encodes a protein that together with proteins encoded by Rad1 and Rad9 form the 9-1-1 checkpoint protein complex. This complex plays a central role in the DNA damage-induced checkpoint response. Involved in G2/M transition of mitotic cell cycle, meiotic and mitotic nuclear division. | https://flybase.org/reports/FBgn0026417 |

We performed a protein network analysis on the entire gene list using STRING (Szklarczyk et al. 2019). The full STRING protein interaction results can be found in Supplementary File 1. Many protein products of genes in autosomal QTL regions have direct interactions, and four proteins with functions related to reproduction (bold in Table 5) interact directly in an insular protein network: dynein light chain Tctex-type, intra-flagellar transport protein 74 homolog, tubulin-specific chaperone D, and cytoplasmic dynein 2 heavy chain 1.

## Discussion

One or more loci on the X chromosomes of both species have a large impact on hybrid male sterility in the *An. coluzzii* × *An. quadriannulatus* hybrids. In both multiple imputation QTL models presented (with and without X by 2L epistasis) the cumulative effect of the X chromosome accounted for greater than 60 % of the phenotypic variation. These results support the theory of large X effect on hybrid male sterility due to its hemizygosity in the heterogametic sex (*Anopheles* males) where recessive sterility genes are expressed, but cannot discount the role of faster X evolution, which we did not test. The large proportion of variance explained by the X in our *An. coluzzii* × *An. quadriannulatus* backcross contrasts with the *An. coluzzii* × *An. arabiensis* backcross (to *An. arabiensis*) performed by Slotman et al. (2004). Slotman et al. (2004) found that the X chromosome accounted for only 4.7% of the variance in the hybrid male sterility phenotype while each of five autosomal QTL accounted for 3.4% to 20.3% of the variance (Table 4, Slotman et al., 2004). However, the relatively small proportion of the variance explained by the X in the *An. coluzzii* × *An. arabiensis* backcross resulted, at least in part, from the inviability observed when hybrids inherited the *An. coluzzii* X (only 9.7% of hybrid males inherited the *An. coluzzii* X, Table 1 of Slotman et al., 2004).

When *An. coluzzii* × *An. arabiensis* F1 hybrids were backcrossed to *An. coluzzii,* the X accounted for 39.5% of the variance and additive effects of autosomal QTL accounted for 4.2-7.9% of the variance (Slotman et al., 2004). The differences in the effect size of the X between crosses, and directions of a back cross between two species (Slotman et al., 2004) is not unexpected. First, the X chromosome has different effects on inviability in each direction of a cross. Second, the genetic architecture of DMIs are unique to each species pair. Additionally, DMIs accumulate over time faster than linearly, and the genetic architecture of individual DMIs becomes more complex as additional mutations accumulate (Matute et al., 2010; Matute and Gavin-Smyth, 2014, Moyle and Nakazato, 2010, Orr and Turelli, 2001).

Unfortunately, due to the *Xag* inversion, the entire X chromosome is effectively one QTL, and we have limited resolution to identify candidate genes on this chromosome. While the 1.5 LOD interval of the X chromosome QTL is 8.53 Mb, the entire span of the QTL is 14.2 Mb, or 58% of the X chromosome. However, we determined which autosomal *An. coluzzii* loci contributed to sterility when introgressed into a predominantly *An. quadriannulatus* genomic background, and vice versa, by dividing the dataset by the parental origin of the X chromosome. This analysis revealed that the epistatic interaction between the 2L and the X QTL is driven by the interaction between *An. quadriannulatus* QTL 2L homozygotes and the hemizygous *An. coluzzii* X chromosome (Figures 3, 4). Individuals with this genotype combination are among the most sterile in the cross (Figure 4), and the 2L QTL is absent in the *An. quadriannulatus* X chromosome CIM results (Figure 3).

The 3L-A QTL sterility QTL in our *An. coluzii* × *An. quadriannulatus* cross overlaps entirely with the 3L QTL identified in the *An. coluzzii × An. arabiensis* cross (backcrossed to *An. arabiensis,* Figure 2.D of Slotman et al., 2004). No *An. arabiensis* homozygotes were sampled in this region when the hybrid inherited the *An. coluzzii* X chromosome. Thus, the *An. coluzzii* X chromosome interacts epistatically with this same region of 3L in both *An.* quadriannulatus and *An. arabiensis* to contribute to hybrid male sterility. We hypothesize that the 3L-A QTL harbors an ancestral allele that is shared between *An.* quadriannulatus and *An. arabiensis* that has not been subjected to introgression between *An. coluzzii* and *An. arabiensis*.

Due to the rapid radiation of the *Anopheles gambiae* species complex, incomplete lineage sorting, and ongoing introgression between member species via fertile F1 hybrid females, the topology of the species complex phylogeny, which is based upon the X chromosome, is discordant with the whole-genome topology (Figure 1 of Fontaine et al., 2015). Bi-directional introgression between *An. arabiensis* and the ancestor of *An. gambiae s.s.* + *An. coluzzii* has reduced the genetic distance between the autosomes of these species relative to the X chromosome. As a result, *An. arabiensis* is sister to *An. quadriannulatus* in the X chromosome phylogeny but is sister to *An. gambiae s.s.* + *An. coluzzii* in the whole genome phylogeny.

While gene tree phylogenies vary across the genome due to the reasons outlines above, on the 3L, where we find overlap of sterility QTL between *An. quadriannulatus* and *An. arabiensis* crosses, the most commonly found gene phylogenies are those that group *An. arabiensis* closer to *An. coluzzii* than *An. quadriannulatus* (Figure 3 of Fontaine et al., 2015). If we assume that genes involved in post-zygotic reproductive isolation between this species trio evolved first on the X chromosome between the ancestor of *An. gambiae s.s.* + *An. coluzzii* and the ancestor of *An. arabiensis* + *An. quadriannulatus*, we can hypothesize that the 3L-A QTL shared between these crosses may have an X chromosome-like topology despite the homogenizing effect of introgression between the *An. coluzzii* and *An. arabiensis* autosomes. This topology is found in ∼12.6% of 50-kb genomics windows on the 3L (Figure 3 of Fontaine et al., 2015; see “other, A+Q trees”).

Our hypothesis that the 3L-A by X chromosome DMI may have been the first to arise between the ancestor of *An. gambiae s.s.* + *An. coluzzii* and the ancestor of *An. arabiensis* + *An. quadriannulatus* can be tested in the following ways. First, this sterility QTL should also exist in a cross between *An. gambiae s.s.* and *An. arabiensis* or *An. quadriannulatus,* and second, it should be absent in a cross between *An. arabiensis* and *An. quadriannulatus*. No hybrid sterility or inviability exists between *An. coluzzii* and its sister species *An. gambiae s.s.*, so we expect that the same loci will be responsible for reproductive isolation between both of these species and either *An. arabiensis* or *An. quadriannulatus*. A cross between *An. arabiensis* and *An. quadriannulatus* may yield more interesting results; it would help us to understand the order by which individual DMI arose between these species pairs, and how this effects our understanding of the “snowball effect” of DMI evolution in the context of rapidly radiating species. For example, while the 3L-A by X chromosome DMI has been maintained between *An. coluzzii* and both *An. arabiensis* and *An. quadriannulatus*, additional DMI may have evolved in a “snowball effect” type scenario between *An. coluzzii* and *An. quadriannulatus,* where there is no evidence historical autosomal introgression (Fontaine et al., 2015), whereas DMI may have been lost as a result of autosomal introgression between *An. coluzzii* and *An. arabiensis*, or their relative rate of evolution may have been slower.

Our candidate sterility gene analysis identified seven genes (out of 349) within autosomal *An. coluzzii × An. quadriannulatus* hybrid male sterility QTL that had protein functions related to tubulin or microtubule biogenesis, structure, or function. Four of these (AGAP004782, AGAP001219, AGAP010790, and AGAP001229) have experimentally determined, direct interactions with each other, while another interaction is predicted due to co-expression (AGAP004724 and AGAP010790) (Table 5, Supplementary File 1). While these genes are found within autosomal QTL that contribute to a small proportion of variation in the sterility phenotype, one or more could have an interaction with an X chromosome locus that is disrupted during hybrid testes morphogenesis and/or spermatogenesis. In *Drosophila,* spermatogenesis progresses through mitosis of germ cells, meiosis, and finally the differentiation of spermatids into mature sperm (White-Cooper et al., 2009). Infertility in *Drosophila* hybrid males is associated with spermiogenic (post-rather than pre-meiotic) failure associated with sperm individualization and maturation (Wu et al., 1992). Recent work in the *Anopheles gambiae* complex demonstrates that hybrid sterility results in part from pre-meiotic arrest in degenerate testes (Liang et al, 2019).

Sperm flagellum elongate from the spermatid head after meiosis. The axoneme is the central component of the sperm flagellum and has a 9+2 structure of a central pair of microtubules surrounded by nine doublet microtubules. Doublet microtubules have an inner and outer dynein arm that serve to attach and detach to neighboring doublet microtubules, and function in the motor activity of the flagellum. The axoneme is surrounded by a mitochondrial sheath that serves to provide ATP and power motor activity. Tektins are located near the junction points of dynein arms and neighboring microtubules (Gagon, 1995; Linck et al., 2016). Seven of the ten reproduction genes have PANTHER protein functions related to tubulin or microtubule biogenesis, structure, or function: gamma-tubulin complex component 5, microtubule-associated protein futsch, tubulin-like protein alpha-4B-related, dynein light chain Tctex-type, intra-flagellar transport protein 74 homolog, tubulin-specific chaperone D, and cytoplasmic dynein 2 heavy chain 1. The latter five genes (bold, Table 5) in this list have direct interactions in the STRING protein interaction network, and may represent a candidate genetic pathway whose disruption results in hybrid male sterility in the *An. coluzzii × An. quadriannulatus* cross. Interestingly, another tubulin-specific chaperone, E-like, is required for sperm individualization and male fertility in *Drosophila* (Nuwal et al., 2012).

The three genes without tubulin/microtubule functions are: transcription factor ken, protein lingerer, and a checkpoint protein Hus1-like. Mutations in the ken and barbie transcription factor result in malformed genitalia in *Drosophila* (Lukacsovich et al., 2003). lingerer encodes a putative RNA binding protein and *Drosophila* lingerer mutant males fail to withdraw their genitalia after copulation (Kuniyoshi et al., 2002). Finally, Hus1-like is a component of the 9-1-1 checkpoint protein complex, which coordinates DNA damage sensing, cell cycle progression, and DNA repair (Cotta-Ramusino et al., 2010). Future analyses of the molecular evolution of these genes and their regulatory regions between member species of the *An. gambiae* complex may yield insight into the specific alleles responsible for hybrid male sterility between *An. coluzzii*, *An. quadriannulatus*, and *An. arabiensis*.

In this study we have determined that the X chromosome accounts for that majority of the variance in the hybrid male sterility phenotype in a cross between *An. coluzzii* and *An. quadriannulatus*. We have also narrowly defined autosomal QTL which have a statistically significant contribution to sterility despite their smaller effect sizes in comparison to the X. The shared, autosomal sterility QTL between the *An. coluzzii × An. quadriannulatus* and *An. coluzzii × An. arabiensis* backcrosses suggest that these may represent important autosomal regions involved in the post-zygotic reproductive isolation of *An. coluzzii* from its sister species that diverged early in the radiation of the *An. gambiae* species complex. Future analyses into factors contributing to female hybrid sterility in the *An. coluzzii × An. quadriannulatus* cross will add to our knowledge of the genetics of speciation in this evolutionarily and medically important group of human malaria mosquitoes.

## Conflict of Interest

The authors declare that the research was conducted in the absence of any commercial or financial relationships that could be construed as a potential conflict of interest.

## Author Contributions

K.C. Deitz, W. Takken and M.A. Slotman designed the experiment. K.C. Deitz and M.A. Slotman, performed the experiment. K.C. Deitz analysed the data. K.C. Deitz and M.A. Slotman wrote the manuscript. W Takken edited the manuscript.

## Funding

This study was funded by a National Science Foundation Doctoral Dissertation Improvement Grant (award # 1601675) to K.C. Deitz and M.A. Slotman, a Texas A&M University Genomics Seed Grant to K.C. Deitz and M.A. Slotman, and a Texas A&M University Dissertation Fellowship to K.C. Deitz.

## Acknowledgments

We thank Patrick Reilly and Andrew Taverner for helpful discussions regarding the analysis of the dataset and Leon Westerd for assistance with mosquito rearing. We are also thankful to BEI Resources for providing mosquito strains.

## Supplementary Material

Supplementary tables can be found in Deitz_et_al_2020_Supplementary_Tables.docx.

## Data Availability Statement

Raw, whole-genome sequencing data of mosquito colonies used in this study to generate pseudo-references are available on the NCBI Sequence Read Archive at project accession number PRJNAXXXXX (data will be uploaded upon acceptance of the manuscript). Genotype data for hybrids can be accessed on Dryad (https://doi.org/XX.XXXX/dryad.XXXXX) (data will be uploaded upon acceptance of the manuscript).

---

**Table S1.**
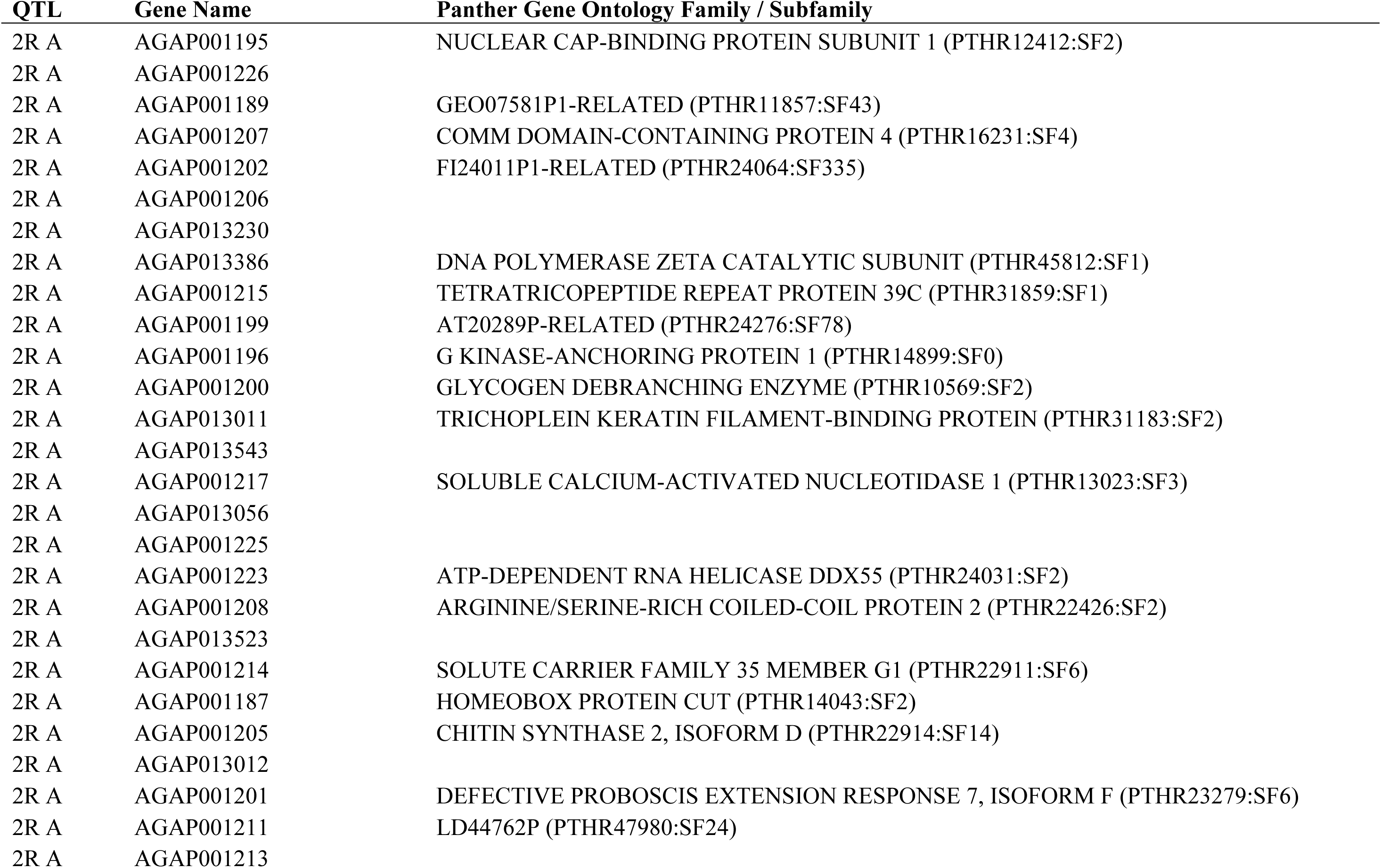
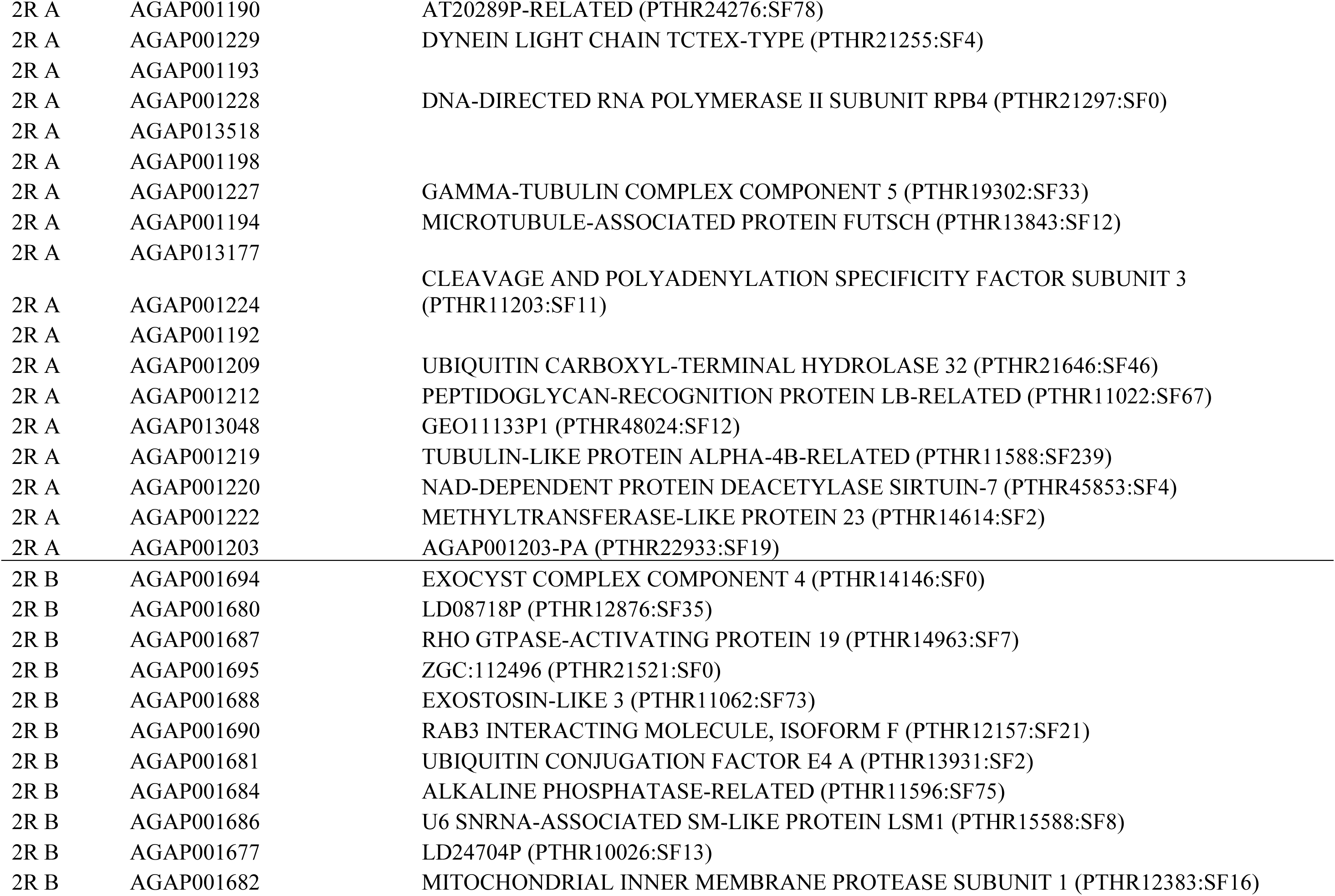
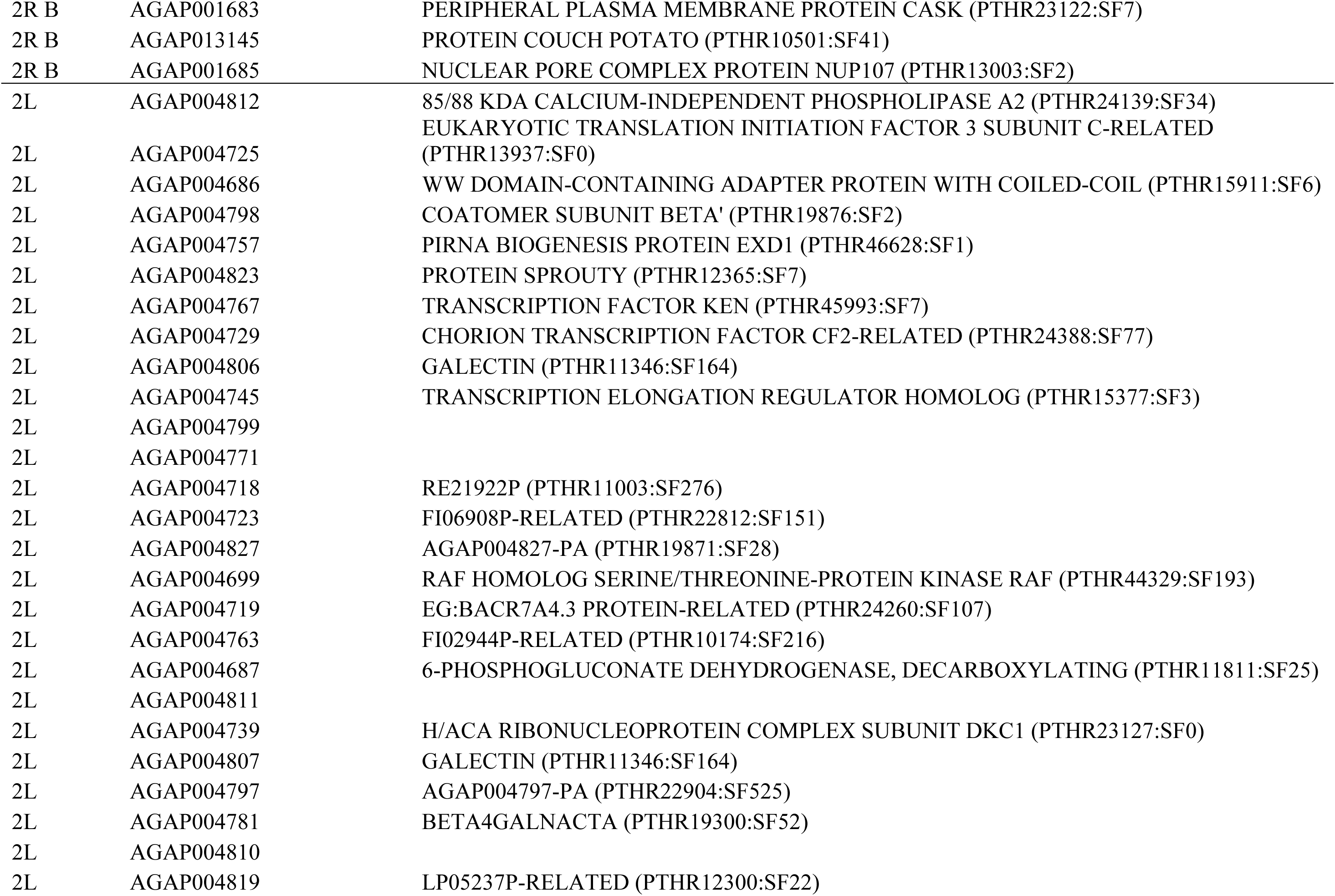
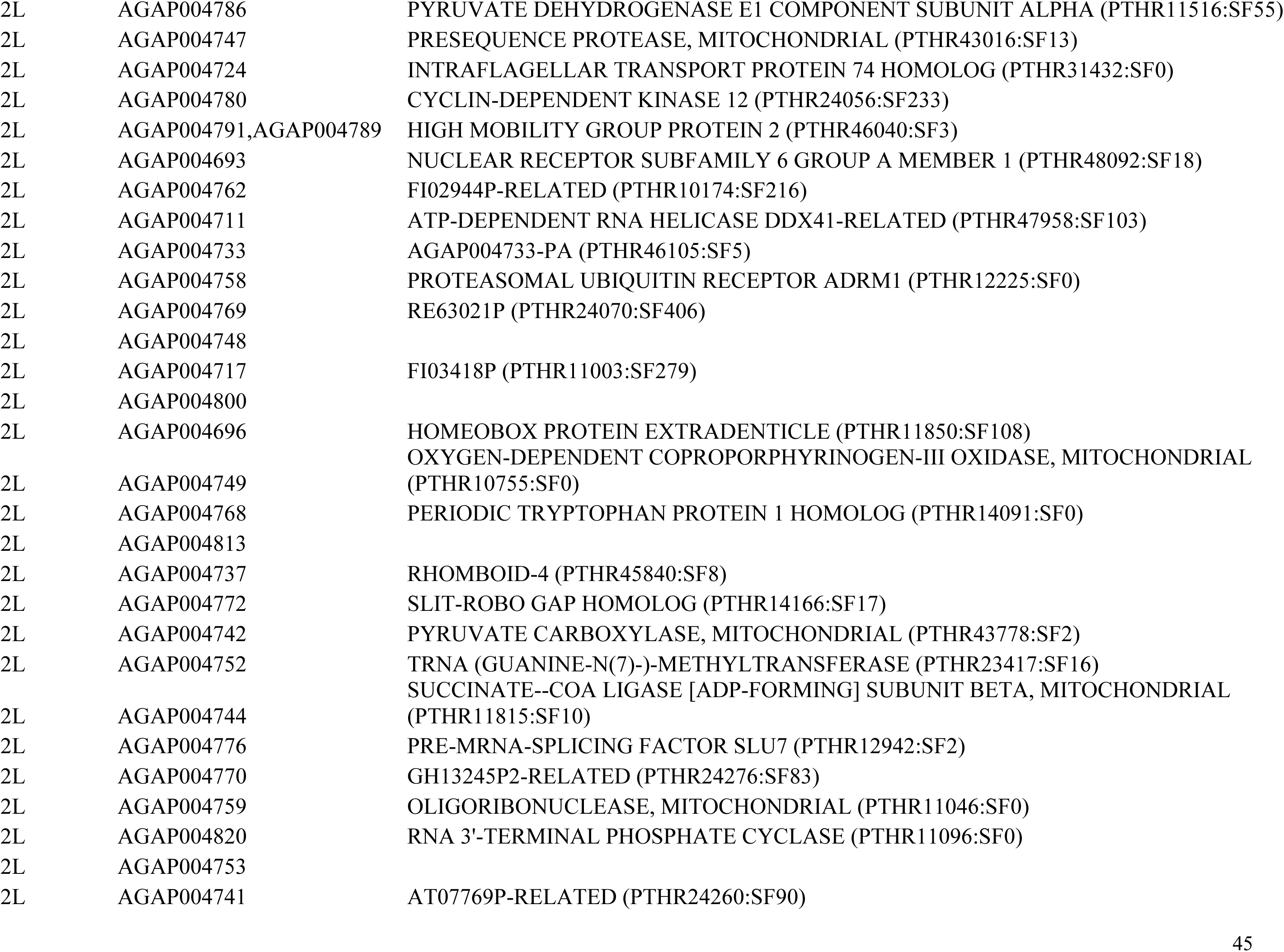
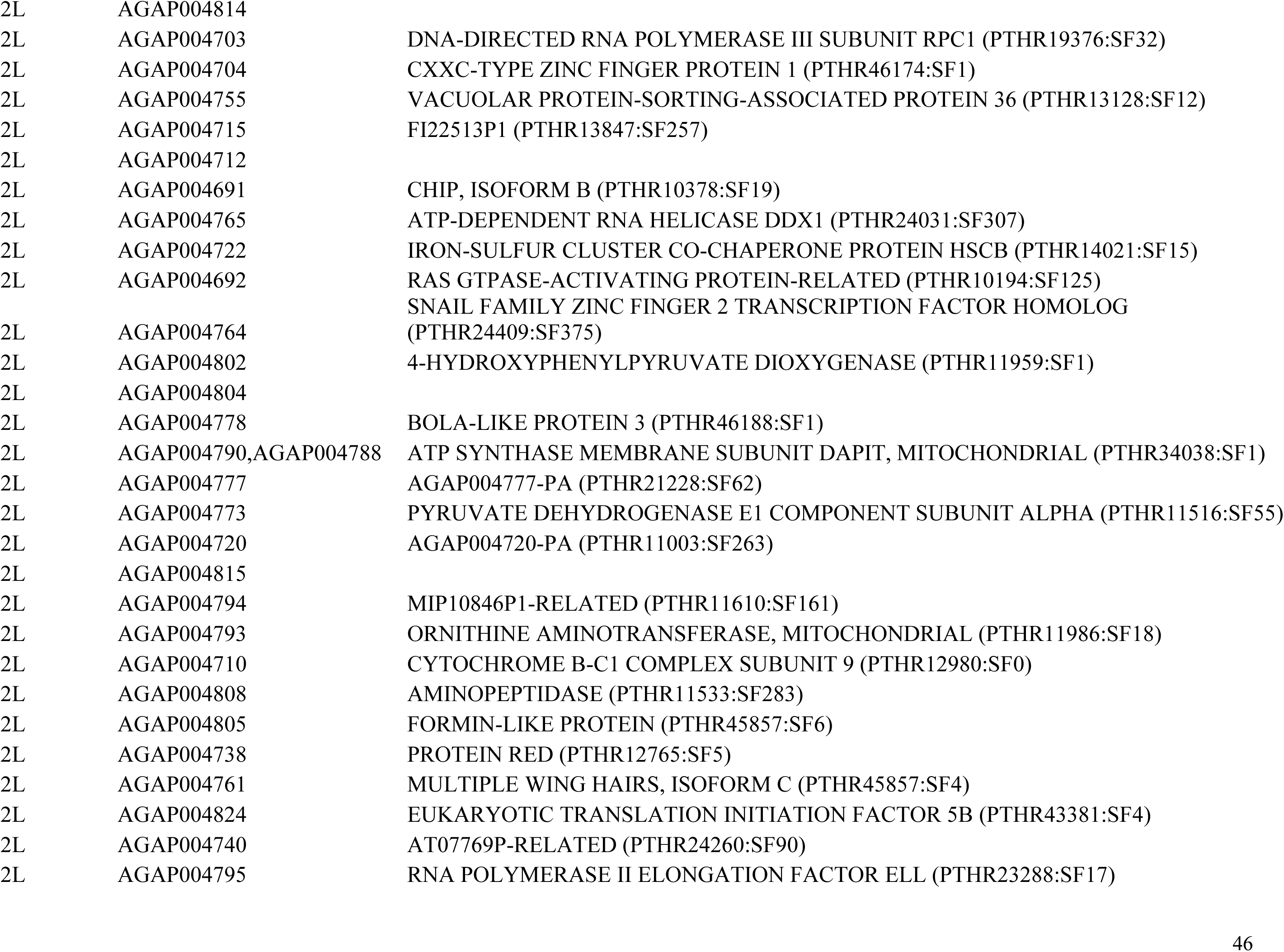
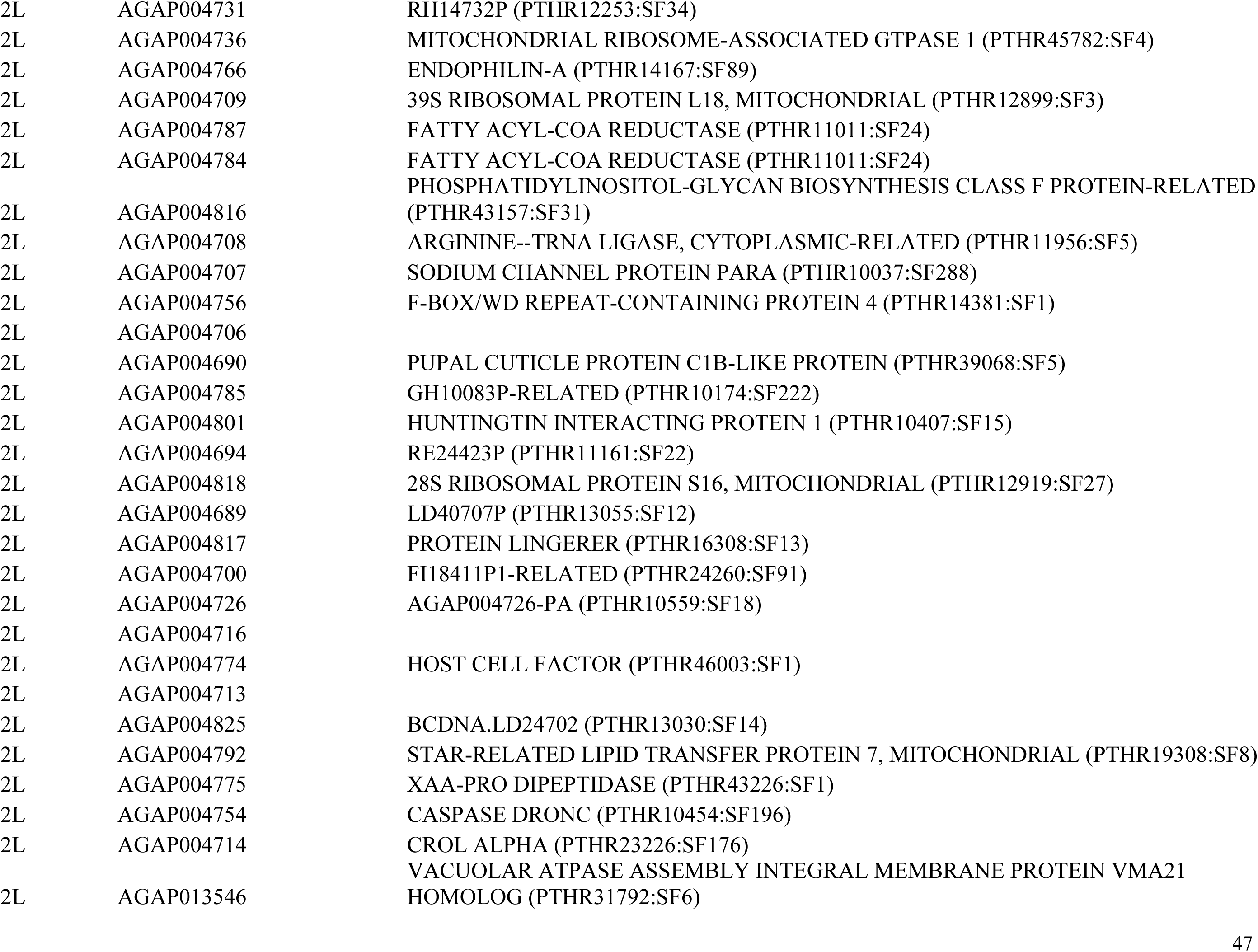
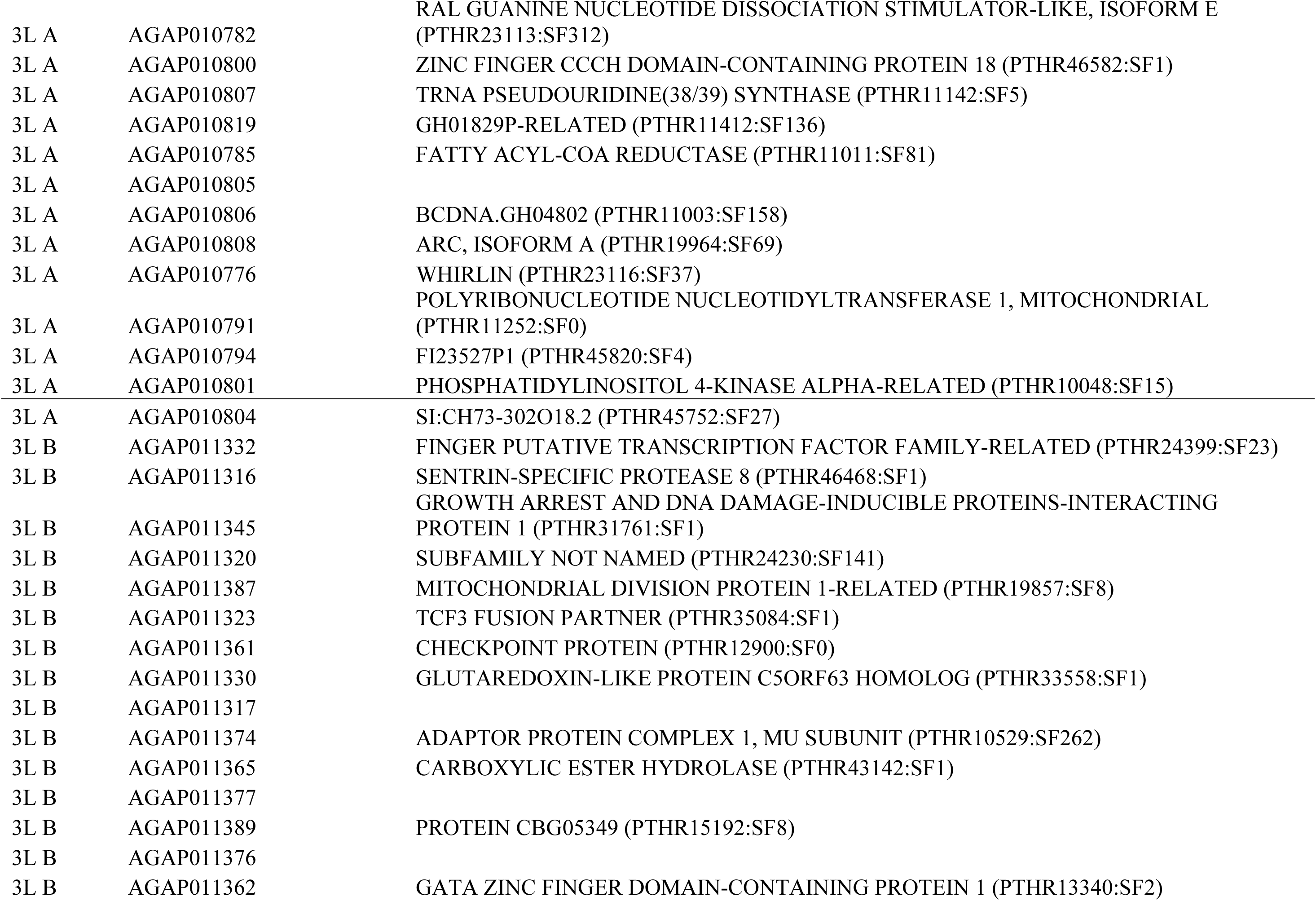
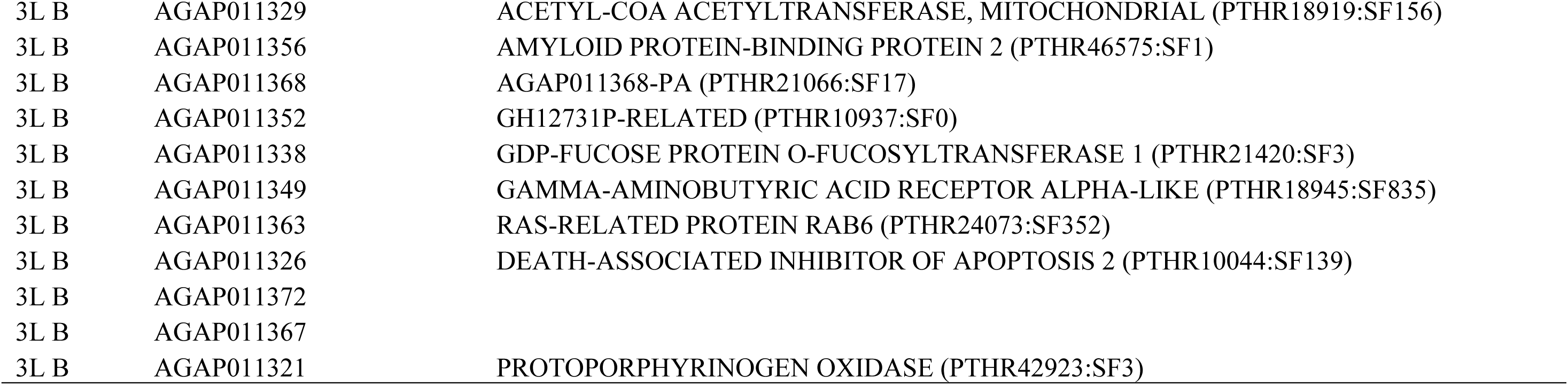
The complete list of annotated genes within autosomal QTL and their PANTHER gene ontology families / sub-families (if available).

## Notes

### Competing Interest Statement

The authors have declared no competing interest.

### Summary of Updates

Section on hybrid male inviability revised. All figures updated.

